# Generating real-world tumor burden endpoints from electronic health record data: Comparison of RECIST, radiology-anchored, and clinician-anchored approaches for abstracting real-world progression in non-small cell lung cancer

**DOI:** 10.1101/504878

**Authors:** Sandra D. Griffith, Melisa Tucker, Bryan Bowser, Geoffrey Calkins, Che-hsu (Joe) Chang, Ellie Guardino, Sean Khozin, Josh Kraut, Rebecca Miksad, Paul You, Deb Schrag, Amy P. Abernethy

**Author notes:** Corresponding author (AA).

## Abstract

Real-world evidence derived from electronic health records (EHRs) is increasingly recognized as a supplement to evidence generated from traditional clinical trials. In oncology, tumor-based Response Evaluation Criteria in Solid Tumors (RECIST) endpoints are collected in clinical trials. The best approach for collecting similar endpoints from EHRs remains unknown. We evaluated the feasibility of a traditional RECIST-based methodology to assess EHR-derived real-world progression (rwP) and explored non-RECIST-based approaches. In this retrospective study, cohorts were randomly selected from Flatiron Health’s database of patient-level EHR data in advanced non-small cell lung cancer. A RECIST-based approach was tested for feasibility (N=26). Three non-RECIST abstraction approaches were tested for feasibility, reliability, and validity (N=200): (1) radiology-anchored, (2) clinician-anchored, and (3) combined. RECIST-based cancer progression could be ascertained from the EHRs of 23% of patients (6/26). In 87% of patients (173/200), at least one rwP event was identified using both the radiology- and clinician-anchored approaches. rwP dates matched 90% of the time. In 72% of patients (124/173), the first clinician-anchored rwP event was accompanied by a downstream event (e.g., treatment change); the association was slightly lower for the radiology-anchored approach (67%; 121/180). Median overall survival (OS) was 17 months (95% confidence interval [CI]: 14, 19). Median real-world progression-free survival (rwPFS) was 5.5 (95% CI: 4.6, 6.3) and 4.9 months (95% CI: 4.2, 5.6) for clinician-anchored and radiology-anchored approaches, respectively. Correlations between rwPFS and OS were similar across approaches (Spearman’s rho: 0.65-0.66). Abstractors preferred the clinician-anchored approach as it provided more comprehensive context. RECIST cannot adequately assess cancer progression in EHR-derived data due to missing data and lack of clarity in radiology reports. We found a clinician-anchored approach supported by radiology report data to be the optimal, and most practical, method for characterizing tumor-based endpoints from EHR-sourced data.

## Introduction

In parallel with expanding treatment options for cancer patients and new scientific paradigms, clinical trials have become increasingly complex. However, patient outcomes remain the benchmark by which new cancer interventions are measured. The efficacy measurement of an intervention in a clinical trial, however, may not directly translate into effectiveness measured in real-world settings [1]. Real-world evidence (RWE) sources, including data derived from electronic health records (EHRs), are a valuable resource to understand the experience of cancer patients treated in the real world to augment insights from traditional clinical trials [2, 3]. RWE can be leveraged to improve therapy development programs by increasing the external validity of evidence available to support point-of-care treatment decisions [4, 5]. The key to fully unlocking the value of the EHR for these purposes is reliable and obtainable outcome metrics; however, it is unknown whether traditional cancer endpoints other than mortality can be gleaned from EHR data. In particular, we need to determine whether we can apply traditionally defined clinical trial tumor endpoints to EHR data, or whether new definitions of endpoints are needed.

Endpoints based on changes in tumor size are often utilized in solid tumor clinical trials to provide measures of treatment efficacy [6, 7]. Radiographic images are typically evaluated using the Response Evaluation Criteria in Solid Tumors (RECIST) and are often reviewed by an independent central committee to improve objectivity of tumor assessments [8, 9]. RECIST uses radiographic evidence about changes in the size of designated tumors over time (“target lesions”) combined with the presence or absence of new tumors to map to categories describing the patient’s disease status: “tumor response,” “stable disease,” or “tumor progression.”

Routine clinical practice follows a similar thread of logic for assessment of response to therapy. Dynamics of tumor burden are often grouped into three main categories: “improved” (i.e., “response”), “no change” (i.e., “stable disease”), or “worse” (i.e., “tumor progression”). Determination of these outcomes is based on periodic evaluation of a variety of clinical parameters assessed with a range of methods (e.g., radiology, physical exam, biomarkers, pathology specimen, patient-reported concerns) and summarized in the clinical notes. Depending on the clinical context, EHR documentation of changes in one or more of these parameters may reflect the outcome of an intervention.

Outcomes data within an EHR are likely composite interpretations of information on the dynamics of tumor burden, quality of life metrics, and other quantitative and qualitative assessment and test results. The richness of EHR data presents significant opportunities to develop outcome metrics specific to real-world data but may also add challenges to the determination and interpretation of real-world outcomes. To further complicate outcome determination in an EHR, the relevant data may reside within sources that are structured (i.e., data that are highly organized, such as white blood cell count) or unstructured (e.g., free text clinical notes or PDFs of radiology or biomarker reports). In other words, the EHR holds data about the effectiveness of an intervention but does so in a manner different than a clinical trial dataset. We may need to make sense of the available EHR data in a different way, likely culling it through manual data curation by trained abstractors following precise, pre-defined policies and procedures, through computer-based algorithms mimicking this approach or a combination of both.

Among these evidence sources available in the EHR, which is best to anchor cancer outcomes against? Radiology reports may not consistently include declarative assessments of response that fit in the RECIST framework. On the other hand, clinician assessments (e.g., radiology “overall impressions” or oncology note “assessment and plans”), findings (e.g., physical exams) and results (e.g., tumor markers) are often documented in the EHR, especially when they inform treatment decisions. Therefore, using clinician notes may serve as an alternative or as a supplement to radiology reports to curate cancer outcomes from EHR data.

Our purpose was to identify a practical and efficient method for large-scale abstraction of data for estimation of cancer outcomes from the EHR. Due to the variety of formats and locations in which documentation of outcomes assessments may exist in the EHR, the abstraction approach used to accomplish this goal must: (1) be applicable across multiple EHR systems; (2) be amenable to manual and/or electronic abstraction from unstructured documents; (3) accommodate the nuances of clinical judgement; (4) be reliable despite the possibility of missing data points; and, (5) be efficient to support scaling in order to enable endpoint assessments for large cohorts.

We anticipated that traditional clinical trial approaches to collecting endpoints, such as cancer progression, may need to be modified for the unique features of EHR data. We tested this hypothesis in a preliminary experiment in a small cohort of patients with advanced non-small cell lung cancer (aNSCLC). We then compared several alternative approaches for abstraction of cancer progression events from the EHR in a larger cohort of aNSCLC patients.

## Methods

We conducted experiments to answer the following research questions: (1) can RECIST criteria be feasibly applied to EHR data to assess cancer progression, and (2) what are potential alternate abstraction approaches and how do they perform?

### Data sources

The overall cohort was selected from Flatiron Health’s longitudinal EHR-derived database, which, at the time of this study, included over 210 cancer clinics representing more than 1.2 million active patients across the United States. We identified more than 120,000 and 25,000 patients diagnosed with lung cancer and aNSCLC, respectively. The majority of patients were treated in the community oncology setting. The clinics included in this study used a variety of EHR systems. Demographic, clinical, and outcomes data were extracted out of the source EHR, which included structured data and unstructured documents. To create the database, we aggregated, normalized, and harmonized patientlevel data. Data were processed centrally and stored in a secure format and environment. Structured data (e.g., treatments, labs, diagnosis codes) were mapped to standard ontologies. Dates of death were obtained from a composite mortality variable comprised of the EHR structured data linked to commercial mortality data and the Social Security Death Index [10]. Unstructured data (e.g., clinician notes, radiology reports, death notices) were extracted from EHR-based digital documents via “technology-enabled” chart abstraction [11]. Every data point sourced from unstructured documents was manually reviewed by trained chart abstractors (clinical oncology nurses and tumor registrars, with oversight from medical oncologists). Quality control was conducted for this dataset, including duplicate chart abstraction of a sample of abstracted variables as well as logic checks based on clinical and data considerations.

### Availability of data and materials

The data that support the findings of this study are available from Flatiron Health, but restrictions apply to the availability of these data, which are subject to the de-identification requirements of the Health Insurance Portability and Accountability Act of 1996 (HIPAA) and implementing regulations, as amended. Practically speaking, in order to share select data and data elements, it is necessary to first define the methods of storage, transmission, access rights, and the scope of intended use prior to making any such data available, and an agreement memorializing the same and applicable re-identification restrictions are required for the purposes of ensuring compliance with the data license, de-identification, data protection specifications, and requirements under HIPAA. Please refer any questions or requests regarding data used in this manuscript to Dr. Amy Abernethy.

### Study design and data collection

This retrospective observational study evaluated methods to assess cancer progression from EHR data through two objectives (Fig 1): (1) to evaluate the feasibility of a RECIST approach (Experiment 1) and (2) if a RECIST approach is not feasible, to evaluate three alternative non-RECIST abstraction approaches (Experiment 2).

**Fig 1. Using the EHR to generate a cancer progression endpoint.**

Inclusion criteria for both experiments were: (1) NSCLC patients diagnosed with advanced disease between January 1, 2011 and April 1, 2016; (2) at least two clinical visits on or after January 1, 2011 documented in the EHR; and (3) documentation of initiation of at least two lines of systemic therapy after advanced diagnosis. Advanced disease was defined as a diagnosis of Stage IIIB or metastatic stage IV, or recurrent early disease.

To assess the feasibility of the RECIST approach (Experiment 1), 26 patients were randomly selected from those meeting the inclusion criteria. The sample size was chosen to achieve expected theme saturation consistent with an initial feasibility study [12, 13]. To study the non-RECIST abstraction approaches (Experiment 2), 200 patients were randomly selected from those meeting the inclusion criteria. The sample size was chosen to balance feasibility and the need for a sufficient number of progression and death events to inform descriptive analyses.

### Experiment 1: Feasibility of RECIST criteria

To determine if the data elements required for RECIST version 1.1 [9] evaluation could be abstracted from patient charts, we evaluated the radiology reports in the EHR for the baseline and first treatment response imaging assessment after initiation of first line (1L) systemic therapy. Radiology reports were evaluated for explicit descriptions of “target lesions” as required by RECIST. The subgroup of patients for whom an explicit description of target lesions was not found were also evaluated with a more lenient approach in which abstractors identified target lesions from the available radiology reports. Lesions at least 1 cm in size were classified as measured if their size was numerically documented in the radiology report. The RECIST approach was determined to be potentially feasible for a patient if: (1) their baseline scan was conducted within two months prior to the referenced 1L therapy start date; (2) their initial follow-up scan was performed at least 28 days after the therapy start date; (3) documentation indicated that the patient’s follow-up scan was compared to the baseline scan; (4) documentation existed for lesion measurements on both the baseline and follow-up scans; and (5) all measured and non-measured lesions were specifically described in each imaging report. Although RECIST can be applied to multiple imaging modalities (e.g., CT, MRI, etc.), CT is preferred in oncology for evaluation of the lung [9]. Therefore, in the context of this experiment, we focused on CT and PET/CT including the chest when evaluating radiology reports. Additional evaluation for specific RECIST version 1.1 criteria, such as measurement of longest diameter for tumors and short-axis for lymph nodes, were excluded from this feasibility experiment with the intention of adding such criteria if RECIST were found to be feasible.

### Experiment 2: Comparison of different abstraction methods for assessing real-world cancer progression

Three approaches for determining real-world cancer progression were used to abstract outcomes data from each patient’s chart: (1) A radiology-anchored approach; (2) a clinician-anchored approach; and (3) a combination of both. The radiology-anchored approach is conceptually similar to RECIST in that it focuses on changes in tumor burden observed by imaging. In contrast to RECIST, the radiology-anchored approach is optimized for the type of radiological evidence sources routinely found in EHR data (e.g., impression section within radiology reports). Reports from any imaging modality were considered. The clinician-anchored approach was conceptually focused on the clinician as the synthesizer of signals from the entire patient’s chart. The combined approach considers both radiology reports and the clinician’s synthesis: the date of progression is determined by the earliest source document for the event. Definitions of each approach, evidence sources evaluated, and the logic for calculating progression dates were defined according to Table 1.

**Table 1.** Non-RECIST based approaches to determining cancer progression using EHR data (Experiment 2). Three different abstraction approaches for determining real-world cancer progression: (1) radiology-anchored approach, (2) clinician-anchored approach, and (3) combined approach. For each abstraction approach, the approach definitions, source evidence evaluated, and progression date assignment rules are described.

|  | **Radiology-anchored approach<sup>a</sup>** | **Clinician-anchored approach<sup>a</sup>** | **Combined approach<sup>a</sup>** |
| --- | --- | --- | --- |
| **Definition** | Documented in the radiology report as progression based on the radiologist's interpretation of the imaging | Documented in a clinician's note as cancer progression based on the clinician's interpretation of the entire patient chart, including diagnostic procedures and tests | Documented in either the radiology report or clinician note as cancer progression |
| **Primary and corroborating evidence sources** | *Primary:* Radiology reports | *Primary:* Clinician notes<br>*Corroborating:* Radiology reports | *Primary:* Radiology reports and clinician notes |
| **Cancer progression date** | Date of the first radiology report that indicated a progression event | Date of the first radiology report referenced by the clinician assessment when available, or date of clinician note when no corresponding radiology was conducted or documented | The earliest report date of the available sources of documentation of a progression event |

<sup>a</sup>Each approach also considered pathology reports as potential evidence for progression. However, there were no instances of pathology reports preceding radiology reports in the cohort analyzed for this study. In addition, there were no instances of conflicting information
between radiology and pathology reports. For simplicity of presentation, pathology reports were excluded from this description as a potential evidence source.

### Analysis

For Experiment 1, descriptive statistics (frequencies and percentages) were calculated for patients that met the RECIST feasibility criteria described above. Results were summarized for the percentage of cases that met all criteria, under both the strictest definition requiring explicit mention of target lesions, as well as the more lenient definition.

For Experiment 2, demographics, clinical and tumor characteristics, and treatment types were summarized using descriptive statistics (medians and interquartile ranges or frequencies and percentages, as appropriate). Progression events within the first 14 days of the start of 1L therapy were excluded from all calculations as they occurred too early to reflect treatment effectiveness.

We compared the three pre-defined data abstraction approaches to assess cancer progression from RWD as per Fig 1. For each of the three abstraction approaches, the proportion of patients with a progression event were computed with accompanying 95% confidence intervals (CIs). The concordance of progression events and associated dates were assessed across abstraction approaches. Discordant cases were reviewed to determine the source of the discrepancies. Among patients with at least one progression event, we assessed the frequency (95% CIs) of near-term (within 15 days prior and up to 60 days after the progression date) clinically relevant downstream events, defined as death, start of new therapy line (second or subsequent lines), or therapy stop. The time window was selected to allow for short delays in documentation of cancer progression that might occur when clinician notes were the only source of evidence (15 days prior) and a clinically relevant follow-up period for near-term events appropriate for aNSCLC (60 days after).

Time-to-event analyses for PFS and Time to Progression (TTP) based on real-world data (RWD, specifically rwPFS and rwTTP) as well as overall survival (OS) were performed using standard Kaplan-Meier methods [14]. The index date was defined as the date of 1L therapy start, and the event date was defined for each outcome: the date of the first progression event (rwTTP); the first progression event or date of death (rwPFS); or the date of death due to any cause (OS). Patients who were lost to follow-up or had not yet experienced the relevant event by the end of the study period were censored at the date of last confirmed activity (last clinical visit or drug administration date). Medians, 95% CIs, and Kaplan-Meier curves were calculated for each outcome.

To assess the correlation between rwPFS and OS, the cohort was restricted to patients with a documented death event (N=123). For the correlation between rwTTP and OS, the cohort was further restricted to patients with documentation of both progression and death events (N=112 for the clinician anchored approach; 113 for the radiology anchored approach and for the combined approach). Spearman’s rank correlation coefficient (Spearman’s rho) was used for all correlation analyses.

The reliability of progression abstraction was assessed by calculating inter-rater agreement for the first progression event in a random subset of patients (N=55) who were abstracted in duplicate. Of note, patients may have had anywhere from zero to multiple progression events within or across multiple lines of therapy; however, only up to the first progression event was included for the inter-rater agreement analysis. Agreement was calculated at the patient level and considered from multiple perspectives: (1) agreement on the presence or absence of at least one progression event (event agreement); (2) in cases where both abstractors found at least one progression event, agreement on when the progression occurred (date agreement); and, (3) a combined approach where both the absence or presence of a progression event and the date of the event, if one was found, contribute toward agreement (overall agreement). Date and overall agreement were calculated for exact date, as well as for 15-day and 30-day windows between dates.

Finally, after completion of the experiment, open-ended qualitative feedback was collected from abstractors regarding the feasibility, consistency, and scalability of different approaches.

## Results

### Experiment 1: Feasibility of RECIST criteria

In Experiment 1, a cohort of 26 aNSCLC patients was evaluated for the feasibility of the RECIST approach (Fig 2). When using a strict definition of RECIST that required radiologist-defined target lesions, none of the patient charts (0%) yielded data suitable for assessing cancer progression. When more lenient criteria were applied that did not require explicit mention of target lesions, only 15 charts (58%) had radiology reports describing appropriate scans available for any type of RECIST assessment. Of these, only 8 (31% of Experiment 1 cohort) demonstrated evidence of direct comparison of all measured key lesions between two timepoints. Even fewer (6, 23% of Experiment 1 cohort) had evidence that all non-measured key lesions were specifically followed between the two timepoints. Due to the lack of feasibility and very low completeness (0-23%, depending on requirement for explicit documentation of key lesions), endpoints obtained from the RECIST approach were not further analyzed or compared to the other abstraction approaches.

**Fig 2. Assessing applicability of RECIST for defining cancer progression in real-world EHR data in Experiment 1**. 26 patient charts were randomly selected from the overall cohort of 7584 patients with at least 2 clinical visits and 2 lines of therapy (LoT). RECIST criteria were applied and the numbers of patients meeting the various criteria were recorded.

^a^Appropriate scans were defined as a baseline scan (PET/CT or CT chest) within 2 months prior to therapy start AND a scan (PET/CT or CT chest) to follow-up response at least 28 days after therapy start.

^b^To be “directly followed,” radiology reports needed to sequentially describe all documented lesions of >1 cm to track changes in size.

### Experiment 2: Comparison of different abstraction methods for assessing real-world cancer progression

The study cohort for Experiment 2 consisted of 200 patients, for whom demographic and clinical characteristics are described in Table 2. The median age at advanced diagnosis was 66 years, half were women, 63% were stage IV at diagnosis, most had non-squamous cell histology (73%), and most had a history of smoking (85%).

**Table 2.** Demographic and clinical characteristics of Experiment 2 cohort.

| Variable | N=200 |
| --- | --- |
| **Demographics** |  |
| Age at advanced diagnosis (years), median [IQR] | 66 [57;72] |
| Age at advanced diagnosis category: |  |
| <55 years | 37 (19%) |
| 55-64 years | 58 (29%) |
| 65+ | 105 (53%) |
| Gender: |  |
| Female | 100 (50%) |
| Male | 100 (50%) |
| Race/ethnicity: |  |
| White | 137 (69%) |
| Black or African American | 15 (8%) |
| Asian | 6 (3%) |
| Other race | 15 (8%) |
| Unknown/missing | 27 (14%) |
| Region: |  |
| Northeast | 62 (31%) |
| Midwest | 36 (18%) |
| South | 67 (34%) |
| West | 35 (18%) |
| **Clinical characteristics** |  |
| Group stage at diagnosis: |  |
| Stage I | 13 (7%) |
| Stage II | 8 (4%) |
| Stage III | 44 (22%) |
| Stage IV | 125 (63%) |
| Group stage is not reported | 10 (5%) |
| Histology: |  |
| Non-squamous cell carcinoma | 145 (73%) |
| Squamous cell carcinoma | 46 (23%) |
| NSCLC histology NOS | 9 (5%) |
| Smoking status: |  |
| History of smoking | 169 (85%) |
| No history of smoking | 25 (13%) |
| Unknown/not documented | 6 (3%) |
| First-line therapy class: |  |
| Platinum doublets | 102 (58%) |
| Single agent chemotherapies | 28 (16%) |
| Platinum doublet + bevacizumab | 20 (11%) |
| EGFR TKIs | 18 (10%) |
| Single-agent chemotherapy + bevacizumab | 2 (1%) |
| Bevacizumab monotherapy | 2 (1%) |
| Other therapies | 2 (1%) |
| Clinical study drug-based therapies | 1 (0.6%) |
| PD-1/PD-L1-based therapies | 1 (0.6%) |
| Treatment setting: |  |
| Community | 194 (97%) |
| Academic | 6 (3%) |
| Follow-up time from advanced diagnosis (months), Median [IQR] | 13 [9.0;21] |

In the majority of patients (N=173, 87%) both radiology- and clinician-anchored approaches identified at least one progression event per patient. In addition to those events identified by both approaches, 7 patients (4% of the Experiment 2 cohort) had radiology-anchored progression events without confirmation in a clinician note, leading to a total of 180 patients with a radiology-anchored progression event (90%). Of those seven cases without clinician confirmation, the discordant information could be attributed to one of two reasons: (1) the radiology report was the last available document in the chart and the clinician assessment did not exist or was unavailable in the chart (5/7), signifying possible patient death, pending clinical visit, referral to hospice, or loss to follow-up, or (2) there was disagreement between the radiology report and the assessment in the clinician note (2/7). In both of these discordant cases, the radiology report recorded progression, but the clinician determined stable disease

For the 173 patients where a progression event was identified by both the radiology- and clinician-anchored approaches, progression dates matched between the two approaches for 156 patients (90%). Among the 17 patients (10%) with differing progression event dates, the radiology-anchored approach identified an earlier progression date in almost all cases (N=16, 93%). In all 16 cases, the clinician made a determination of stable disease immediately following the radiology-anchored progression event but subsequently documented a progression event. In the one final discordant case, the clinician-anchored progression event did not have a corresponding scan documented in the chart.

The association between cancer progression and near-term downstream events (15 days prior up to 60 days after), including treatment stop, treatment change, or death, is displayed in Table 3. Of the 173 patients with a cancer progression identified by the clinician-anchored approach, 124 (71.7%) experienced a near-term downstream event accompanying their first cancer progression. Of the 180 patients with a cancer progression identified by the radiology-anchored approach, 121 (67.2%) experienced a near-term downstream event accompanying their first cancer progression. The combined approach using the first clinician- or radiology-anchored event showed identical results to the radiology-anchored approach.

**Table 3.** Likelihood of predicting downstream events in Experiment 2.

|  | **Radiology-anchored approach** | **Clinician-anchored approach** | **Combined approach** |
| --- | --- | --- | --- |
| Number of patients | 180 (90%) | 173 (87%) | 180 (90%) |
| with at least one progression event (% of Experiment 2 cohort) |  |  |  |
| Number of patients with a downstream event<sup>a</sup> | 121 | 124 | 121 |
| % of patients with an associated downstream event (95% CI) | 67% (60, 74) | 72% (65, 79) | 67% (60, 74) |

<sup>a</sup> Clinically relevant downstream events defined as death, start of new therapy line (second or subsequent lines), or therapy stop. Downstream events occurred 15 days prior and up to 60 days after the progression date.

Kaplan-Meier curves for time-to-event endpoints (rwPFS and rwTTP) based on each abstraction approach, accompanied by OS, are displayed in Fig 3. Clinician-anchored median rwPFS (Table 4) was higher than its radiology-anchored counterpart (5.5 months [95% CIs: 4.6, 6.3] vs. 4.9 months [95% CIs: 4.2, 5.6]). A similar pattern was observed for rwTTP. Median OS for all patients was 17 months (95% CI 14, 19). Correlations between rwPFS or rwTTP and OS (Table 4) were very similar across abstraction approaches and slightly higher for rwTTP (Spearman’s rho: 0.70; 95% CI: 0.59, 0.78) than for rwPFS (Spearman’s rho [95% CI] ranged from 0.65 [0.53, 0.74] to 0.66 [0.55, 0.75], depending on approach).

**Fig 3. rwPFS, rwTTP and OS in Experiment 2**. Kaplan-Meier survival curves for progression-free survival (PFS) or time to progression (TTP) and overall survival (OS) for all three non-RECIST abstraction approaches.

**Table 4.** Correlation between rwPFS or rwTTP and OS in Experiment 2.

|  | Radiology-anchored approach | Clinician-anchored approach | Combined approach |
| --- | --- | --- | --- |
| Median rwPFS, months (95% CI) | 4.9 (4.2, 5.6) | 5.5 (4.6, 6.3) | 4.9 (4.2, 5.6) |
| rwPFS correlation with OS (95% CI)<sup>a,b</sup> | 0.65 (0.53, 0.74) | 0.66 (0.55, 0.75) | 0.65 (0.53, 0.74) |
| Median rwTTP, months (95% CI) | 5.0 (4.2, 6.1) | 5.6 (4.8, 6.5) | 5.0 (4.2, 6.1) |
| rwTTP correlation with OS (95% CI)<sup>a,c</sup> | 0.70 (0.59, 0.78) | 0.70 (0.59, 0.78) | 0.70 (0.59, 0.78) |

<sup>a</sup>Spearman's rho.
<sup>b</sup>Includes only patients with an observed death (N=123).
<sup>c</sup>Includes only patients with an observed death and a cancer progression event preceding death (N=112 for the clinician-anchored approach; N=113 for the radiology-anchored and the combined approach).

Similar inter-rater agreement was observed across abstraction approaches (Table 5). In 96% to 98% of cases, depending on approach, abstractors agreed on the presence or absence of a first progression event. Overall agreement was also similar for all approaches, ranging from 71% to 73% when 30-day windows between progression event dates were allowed. When considering only cases where both abstractors agreed that at least one progression event occurred (N=48, 49), the progression event dates were within 30 days of each other in 69% to 71% of patients.

**Table 5.** Inter-rater reliability in Experiment 2. Patient charts were abstracted in duplicate by different abstractors and agreement (95% CIs) is reported. Event agreement is based on the presence or absence of at least one cancer progression event. Date agreement is based on when the progression occurred, and only calculated in cases where both abstractors recorded a cancer progression. Overall agreement is based on a combined approach where both the absence or presence of a progression event and the date of the event, if one was found, contribute toward agreement.

| Approach | Patients considered | N | Exact agreement | 15-day window | 30-day window |
| --- | --- | --- | --- | --- | --- |
| Radiology-anchored approach (95% CI) | Event agreement | 55 | 98% (94%, 100%) | - | - |
|  | Date agreement | 49 | 61% (47%, 75%) | 67% (54%, 80%) | 69% (56%, 82%) |
|  | Overall agreement | 55 | 0.64 (0.51, 0.77) | 0.69 (0.57, 0.81) | 71% (59%, 83%) |
| Clinician-anchored approach (95% CI) | Event agreement | 55 | 96% (91%, 100%) | - | - |
|  | Date agreement | 48 | 60% (46%, 74%) | 67% (54%, 80%) | 71% (58%, 84%) |
|  | Overall agreement | 55 | 62% (49%, 75%) | 67% (55%, 79%) | 71% (59%, 83%) |
| Combined Approach (95% CI) | Event agreement | 55 | 98% (94%, 100%) | - | - |
|  | Date agreement | 49 | 61% (47%, 75%) | 69% (56%, 82%) | 71% (58%, 84%) |
|  | Overall agreement | 55 | 64% (51%, 77%) | 71% (59%, 83%) | 73% (61%, 85%) |

Qualitative feedback revealed a strong abstractor preference for the clinician-anchored abstraction approach because it afforded a more comprehensive context. One abstractor reported that the clinician-anchored approach “provide[s] a true comparison of the scan results and guide to overall patient treatment.” Abstractors also reported the clinician-anchored approach to be faster and more straightforward compared to the radiology-anchored approach.

## Discussion

Real-world research based on EHR data requires validated approaches to abstract outcomes in order to glean meaningful insights and ensure that outcomes are consistently described across different locations and studies. What we really want to know about an intervention is if the patient is “improved,” “worse,” or “unchanged.” Many proxies have been proposed to answer this question. In prospective clinical trials, RECIST measurement of tumor size on imaging is a common metric. However, it was unknown if the language of clinical trial endpoints could be directly translated to routine clinical care. In addition, abstraction of outcomes from the EHR needs to facilitate, rather than hinder, research on large, contemporary cohorts. Therefore, we evaluated four approaches, one RECIST-based and three non-RECIST-based, to identify a feasible method suitable for large-scale abstraction of cancer progression from EHRs.

We found that it is not feasible to use standard, or even lenient, RECIST criteria to abstract cancer progression from the EHR. RECIST cannot be applied to EHR-based RWD because outcomes are missing at least 75% of the time. This finding is not surprising given that in the clinical trial setting collection of RECIST data requires enormous resources.

Given the infeasibility of RECIST for ascertaining real-world progression, we tested and compared three alternative approaches to define cancer progression from the EHR using technology-enabled human curation (Table 1). All three non-RECIST approaches for abstraction of routine EHR documentation yielded similar analytic results and identified progression events at a frequency in line with clinical expectations. Progression events were identified in 86 to 90% of cases, regardless of approach. Progression events predicted near-term downstream clinical events, such as treatment change and death, more than two thirds of the time. For those progression events not associated with a near-term clinical event, medical record review confirmed that it was due to reasonable realities of clinical care (e.g., few subsequent treatment options or need for a treatment holiday). Regardless of abstraction approach, the real-world progression variable performed as expected in time-to-event analyses; a median rwPFS of approximately 5 months is similar to that observed in published aNSCLC clinical trials [15, 16]. And, correlations between rwPFS/rwTTP and OS shown here are consistent with, or higher than, results from clinical trials [15, 17, 18]. Given our reliance on RECIST in clinical trials, one might anticipate that the radiology-anchored abstraction approach would be most reliable; however, the clinician-anchored approach was similarly reliable and abstractor feedback indicated that it may be more scalable. This similar reliability between approaches may be due to the overlap between clinician and radiology assessments of disease progression in the real-world setting.

Modest differences in rwPFS for the two methods reflect differences in the underlying conceptual structure. More progression events were identified in the radiology-anchored approach, but the clinician-anchored approach was more likely to be associated with a treatment change or death. This finding suggests that the clinician may be synthesizing information and adjudicating potentially conflicting findings. Median rwPFS was approximately two weeks shorter for the radiology-anchored approach when compared with the clinician-anchored approach, likely because imaging often precedes a clinician visit and/or assessment in real-world settings. However, modest differences and overlapping confidence intervals preclude any strong conclusions. Importantly, no meaningful differences between approaches were observed for the association between rwPFS and OS in this cohort.

Despite these similar quantitative results, abstractor feedback identified substantial differences in the implementation of different approaches. The clinician-anchored approach was favored, as the clinician’s synthesis of radiology and other findings resonated with the broader context of the patient’s clinical journey. In contrast, radiologists may not always have access to the EHR to support interpretation of imaging findings. In addition, abstraction times were reported by abstractors to be shorter for the clinician-anchored approach. These features of the clinician-anchored approach support the feasibility and scalability of abstraction of progression from the EHR. A scalable approach can support research on a large patient population with a rapid turnaround time. This speed accommodates research using contemporary data to answer questions of relevance to current and future patients.

In this study, we identified a clinician-anchored cancer progression abstraction method to be the most practical approach for assessment of tumor-based endpoints derived from information in EHRs, with radiology and pathology reports serving as corroborating evidence. The clinician-anchored approach is based on clinician summaries as the first source of evidence, a concept in line with how EHRs are used by clinicians. When compared with RECIST-based criteria, where even the most lenient interpretation leads to an untenable amount of missing data for most analyses, the clinician-anchored approach yields much more complete endpoint information across patients. The radiology-anchored approach was developed to align conceptually with the perspective of RECIST, directing abstractors to consider radiology reports as the primary source of truth. Based on our results, the radiology-anchored approach is reasonable as well. However, the radiology-anchored approach is potentially less scalable, at least when medical records are sourced from the community oncology setting, where the majority of patients in this cohort received treatment. Further, it is plausible the radiology-anchored approach may not perform as well in situations where determination of progression is more often based on non-radiologic findings: examples include findings manifest on physical exam, symptom history, biomarker evidence, and for care in resource- or access-limited settings.

One might question why we deemed the RECIST-based approach infeasible after only 26 patients. We applied the saturation sampling method used in qualitative research [13, 14]. The goal of our study was to evaluate whether adequately-complete outcomes data could be generated from the EHR applying the rules of RECIST. Even if we set a very lenient feasibility threshold of 75% completeness for abstraction of progression events by RECIST, it became obvious after 20 charts that the results fell far short of this minimum. We continued to enrich the sample to 26 and the poor data completeness persisted.

When measuring cancer progression using the approach outlined here, there are several limitations to consider. First, this proposed approach is susceptible to several sources of subjectivity, including that of radiologists, clinicians, and abstractors. However, the analysis results are encouraging in that they show rwPFS correlates with OS in ways similar to RECIST-based PFS, and the correlation is also similar between the radiology- and clinician-anchored approaches. We implemented clear instructions and trainings to abstractors to reduce this potential source of subjectivity during the abstraction process. Second, although similar inter-rater reliability was observed across approaches, further work, including additional training and abstractor experience with the approach, could improve reliability.

More broadly, there are additional limitations to consider when applying this work across other settings. First, application to other diseases beyond aNSCLC needs to be tested; there may be differences in assessment cadence or availability of biomarker data, as well as fundamental difference in disease course and treatment pace. These potential differences mean that we must develop a set of rules for reconciling conflicting data about progression as well as incorporation of additional data sources. Second, our work predominantly included EHR data generated in the community oncology setting with few academic medical centers. Documentation and treatment patterns may differ across treatment settings, limiting the generalizability of these results beyond the community oncology setting. Third, whether this approach will hold true for other real-world endpoints such as tumor response will require further examination. Lastly, any approach utilizing time-dependent endpoints is susceptible to bias due to data missingness or the frequency of assessments. This is most likely to be the case if assessment frequency is unbalanced across treatment groups and requires further study to characterize the extent of potential bias.

Based on these findings, we foresee two important next steps. In order to fully harness the potential value of EHR data, outcome measures tailored to the unique features of the data source are needed. Once those outcome measures are developed, such as the methodology for abstraction of cancer progression described here, a comprehensive validation framework must be established to enable robust and standardized characterization of such approaches. Second, we need to tailor and expand this outcomes development and validation approach to different contexts and cancer types. It is also important to explore the broader opportunities to assess other outcome metrics suitable for the real-world setting, for example, quality of life. This study lays the foundations upon which to build endpoint measures suited for RWD/RWE research.

## Acknowledgements

The authors would like to thank Rana Elkholi, PhD, and Nicole Lipitz for editorial assistance and Rachael Sorg, MPH, for assistance with statistical analyses.

## References

1. Sox HE, Greenfield S. Comparative Effectiveness Research: A Report from the Institute of Medicine. Ann Intern Med. 2009; 151(3):203–5.

2. Berger ML, Curtis MD, Smith G, Harnett J, Abernethy AP. Opportunities and challenges in leveraging electronic health record data in oncology. Future Oncol. 2016; 12:1261–1274.

3. Khozin S, Blumenthal GM, Pazdur R. Real-world data for clinical evidence generation in oncology. J Natl Cancer Inst. 2017; 109.

4. Martell RE, Sermer D, Getz K, Kaitin KI. Oncology drug development and approval of systemic anticancer therapy by the U.S. Food and Drug Administration. Oncologist. 2013; 18:104–111.

5. Sherman RE, Anderson SA, Dal Pan GJ, Gray GW, Gross T, Hunter NL, et al. Real-world evidence - what is it and what can it tell us? N Engl J Med. 2016; 375:2293–2297.

6. Johnson JR, Williams G, Pazdur R. End points and United States Food and Drug Administration approval of oncology drugs. J Clin Oncol. 2003; 21:1404–1411.

7. Pazdur R. Endpoints for assessing drug activity in clinical trials. Oncologist. 2008; 13 Suppl 2:19–21.

8. Therasse P, Arbuck SG, Eisenhauer EA, Wanders J, Kaplan RS, Rubinstein L, et al. New guidelines to evaluate the response to treatment in solid tumors. European Organization for Research and Treatment of Cancer, National Cancer Institute of the United States, National Cancer Institute of Canada. J Natl Cancer Inst. 2000; 92(3):205–216.

9. Eisenhauer EA, Therasse P, Bogaerts J, Schwartz LH, Sargent D, Ford R, et al. New response evaluation criteria in solid tumours: revised RECIST guideline (version 1.1). Eur J Cancer. 2009; 45:228–247.

10. Curtis MD, Griffith SD, Tucker M, Taylor MD, Capra WB, Carrigan G, et al. Development and validation of a high-quality composite real-world mortality endpoint. Health Serv Res. 2018; doi: 10.1111/1475-6773.12872.

11. Abernethy AP, Gippetti J, Parulkar R, Revol C. Use of electronic health record data for quality reporting. J Oncol Pract. 2017; 13:530–534.

12. Virzi RA. Refining the test phase of usability evaluation: how many subjects is enough? Human Factors: The Journal of the Human Factors and Ergonomics Society. 1992; 34:457–468.

13. Nielsen J, Landauer, TK. A mathematical model of the finding of usability problems. CHI ‘93 Proceedings of the INTERACT ‘93 and CHI ‘93 Conference on Human Factors in Computing Systems. 1993; doi:10.1145/169059.169166.

14. Kaplan EL, Meier P. Nonparametric estimation from incomplete observations. J Am Stat Assoc. 1958; 53: 457–81.

15. Johnson KR, Ringland C, Stokes BJ, Anthony DM, Freemantle N, Irs A, et al. Response rate or time to progression as predictors of survival in trials of metastatic colorectal cancer or non-small-cell lung cancer: a meta-analysis. Lancet Oncol. 2006; 7: 741–6.

16. Laporte S, Squifflet P, Baroux N, Fossella F, Georgoulias V, Pujol JL, et al. Prediction of survival benefits from progression-free survival benefits in advanced non-small-cell lung cancer: evidence from a meta-analysis of 2334 patients from 5 randomised trials. BMJ Open. 2013; doi:10.1136/bmjopen-2012-001802.

17. Yoshino R, Imai H, Mori K, Takei K, Tomizawa M, Kaira K, et al. Surrogate endpoints for overall survival in advanced non-small-cell lung cancer patients with mutations of the epidermal growth factor receptor gene. Mol Clin Oncol. 2014; 2: 731–6.

18. Blumenthal GM, Karuri SW, Zhang H, Zhang L, Khozin S, Kazandjian D, et al. Overall response rate, progression-free survival, and overall survival with targeted and standard therapies in advanced non-small-cell lung cancer: US Food and Drug Administration trial-level and patient-level analyses. J Clin Oncol. 2015; 33: 1008–14.

